# Somatosensory information drives modification of a motor memory

**DOI:** 10.64898/2026.05.11.724318

**Authors:** Hiroki Ohashi, Hayato Nishioka, Yuki Ogasawara, Keitaro Murakami, Kazuhisa Shibata

**Affiliations:** Center for Brain Science, RIKEN, 2-1 Hirosawa, Saitama 351-0106, Japan; SONY Computer Science Laboratories Inc., Tokyo 141-0022, Japan

## Abstract

The role of somatosensory information in motor memory modification remains controversial. One view holds that somatosensory information plays only a supportive role, such as feedback or sensory calibration, and is not directly involved in memory modification itself. An alternative view is that afferent somatosensory information, on its own, can modify motor memory. Here we resolve this controversy by isolating somatosensory information from voluntary motor execution using a hand-exoskeleton robot. Participants first learned a sequential finger movement task through voluntary practice. Their learned finger movements were recorded using a data glove. On subsequent days, they underwent a brief, 30-sec reinstatement session: either exoskeleton-based reinstatement, which reproduced their learned movements without voluntary execution, or voluntary reinstatement, in which they executed the trained task. Exoskeleton-based reinstatement led to subsequent performance gains comparable to those induced by voluntary reinstatement. This gain was not explained by replay of visual information, mental rehearsal, enhanced arousal, or warm-up effects induced by exoskeleton-driven movements. Rather, the results indicate that brief reinstatement of somatosensory information alone is sufficient to drive performance gain. We further found that exoskeleton-based exposure to a sequence that only partially matched the trained sequence yielded performance gains, whereas voluntary execution of the same partially matched sequence did not. This dissociation suggests that somatosensory information broadly activates a relevant motor skill memory, whereas voluntary execution activates memory in a sequence-specific manner. Together, these findings resolve a long-standing controversy in motor learning research by showing that afferent somatosensory information alone enables sequential motor memory modification through a route distinct from voluntary execution.

## Introduction

After initial acquisition, motor skill memories remain modifiable through subsequent experience. This modification of motor memory has typically been studied in the context of voluntary movement, in which self-generated signals, including motor commands and sensory predictions, drive refinement of the motor memory (Shadmehr, Smith, and Krakauer 2010; Wolpert, Ghahramani, and Jordan 1995; Todorov and Jordan 2002; Lotze et al. 2003). Within this framework, somatosensory consequences of movement have been assigned supportive roles for error calculation and sensory calibration and thought to be insufficient, on its own, to drive the modification of a motor skill memory.

However, converging evidence suggests that somatosensory information may play a more direct role in motor memory than these supportive roles imply. Somatosensory representations change with motor learning and are associated with subsequent retention of motor memory (Ostry et al. 2010; Hirano, Sakurada, and Furuya 2020). Disrupting somatosensory cortical processing can impair the retention or consolidation of learned motor skills (Seidler, Noll, and Thiers 2004; Wali 2020; Ebrahimi, van der Voort, and Ostry 2024). Passive movement driven by a robot can facilitate the recall of a previously learned motor memory (Kumar, van Vugt, and Ostry 2021). Together, these findings raise the possibility that afferent somatosensory information may provide a route through which existing motor memories are accessed and potentially modified.

This possibility has remained unresolved because prior approaches have not isolated afferent somatosensory information from other factors that can also modify motor memory. In most motor learning paradigms, somatosensory signals arise as consequences of voluntary movement and are therefore inseparable from motor commands, sensory predictions, attention, and performance-related feedback. Studies with passive movement have provided important evidence that somatosensory information can contribute to memory recall, but these effects have typically been examined as facilitation of memory retrieval rather than motor memory modification. This limitation has left unresolved whether afferent somatosensory information is merely associated with motor memory processes or can itself serve as a driver of the modification of an already acquired motor memory.

To resolve this issue, the present study isolated afferent somatosensory information from voluntary motor execution. On the first day, participants learned a motor skill in a sequential finger movement task through voluntary practice (Censor, Dimyan, and Cohen 2010; Herszage, Sharon, and Censor 2021). On subsequent days, we used a hand-exoskeleton robot (Furuya et al. 2025) to deliver brief, 30-sec reinstatement of the learned finger movements recorded during training on the first day, thereby reinstating skill-related somatosensory information without requiring voluntary movement (exoskeleton-based reinstatement condition). In a voluntary reinstatement condition, participants instead voluntarily executed the same learned finger movements. On the last day, participants performed the sequential finger movement task again. We then tested whether performance gain occurs as a result of the reinstatement sessions.

The brief exoskeleton-based reinstatement resulted in subsequent performance gain to a degree comparable to that induced by the voluntary reinstatement. This gain was also observed when participants engaged in a demanding working memory task during the exoskeleton-based reinstatement, arguing against a primary contribution of explicit mental rehearsal to the performance gain. On the other hand, such gain was not observed when visual display during training on the first day was replayed without finger movements during the reinstatement sessions. These results suggest that brief, 30-sec reinstatement of the afferent somatosensory information is sufficient to induce subsequent performance gain.

Next, we tested whether performance gain induced by exoskeleton-based reinstatement requires a sequence that exactly matches the one used during training. When participants were passively exposed to a sequence partially matched with the trained one during the reinstatement sessions, we found performance gain. This result suggests that exoskeleton-based reinstatement broadly activates relevant motor memories and serves as a trigger that drives the strengthening of those memories. Importantly, the voluntary execution of the same partially-matched sequence did not yield the gain. That is, voluntary reinstatement selectively activates the motor memory in a sequence-specific manner. These results suggest that exoskeleton-based reinstatement and voluntary reinstatement produce performance gain through different neural mechanisms.

Collectively, the present study demonstrated that afferent somatosensory information can, on its own, be sufficient to modify a motor skill memory, resolving the long-standing controversy on the role of somatosensory information in motor memory modification.

## Results

### Reinstatement of somatosensory information is sufficient to strengthen a learned sequential motor skill

In Experiment 1 (see Methods for details), we tested whether reinstating the somatosensation of a learned sequential motor skill, without any voluntary movement, is sufficient to modify the skill.

Participants trained in a four-finger sequential movement task (Karni et al. 1995) with their non-dominant left hand placed on load cells that detected finger taps (Fig. 1a). In each 30-sec trial, they were asked to repeatedly tap a 5-item sequence (trained sequence: 4–1–3–2–4) on a display as quickly as possible in the correct order. Each correct completion of the full sequence was counted as one successful sequence. Following a prior study using the same task and experimental procedure (Herszage, Sharon, and Censor 2021), we used the number of correctly completed sequences per 30-sec trial as the performance measure.

**Fig. 1.**
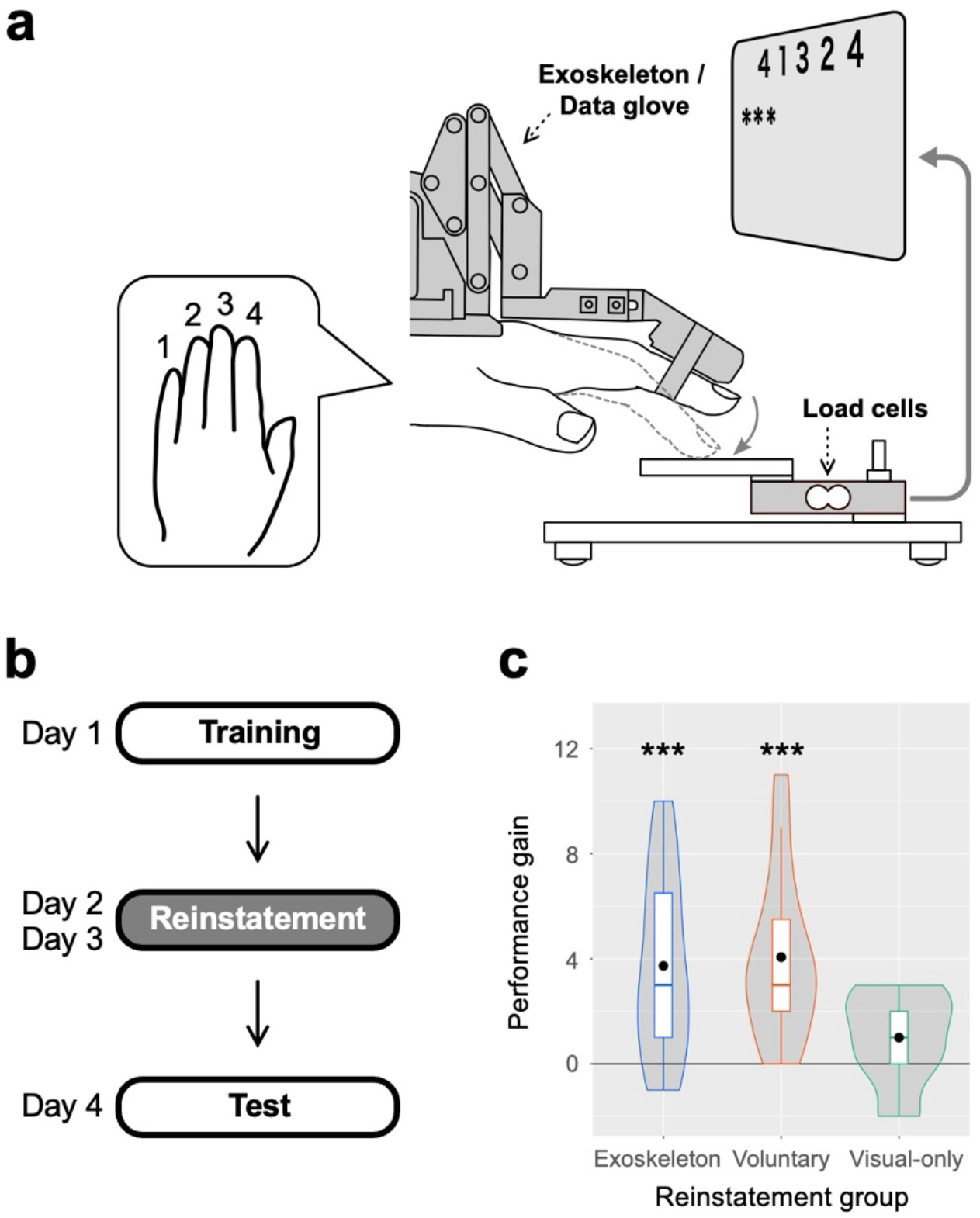
Procedures and results of Experiment 1 **a**, Four-finger sequential movement task. In each 30-sec trial, participants were asked to repeatedly tap a 5-item sequence (4–1–3–2–4) shown on a display as quickly as possible in the correct order with their non-dominant left hand. Each correct completion of the full sequence was counted as one successful sequence. We defined performance as the number of correctly completed sequences per 30-sec trial. In the exoskeleton-based reinstatement group, a hand-exoskeleton robot attached to the left hand reproduced learned finger movement recorded during the training stage for each participant. The same robot was also attached to the left hand of participants in the voluntary and visual-only reinstatement groups. However, no exoskeleton-driven finger movement was generated for these groups. **b**, The complete procedure consisted of one day of training stage, two days of the reinstatement stage, and one day of the test stage. **c**, Performance gain in the test stage, relative to the baseline for the exoskeleton-based (N = 15), voluntary (N = 15), and visual-only (N = 13) reinstatement groups. Box plots are overlaid on violin plots. Black dots indicate mean values across participants. Violin plots show kernel probability densities of individual values. ***P < 0.005.

The complete procedure consisted of one day of the training stage, two days of the reinstatement stage, and one day of the test stage (Fig. 1b), following a prior study (Herszage, Sharon, and Censor 2021). On Day 1 (training stage), all participants completed 12 trials of training. Throughout the stage, participants’ finger movements were recorded by a data glove on their left hand. Performance in the last training trial served as the baseline to test performance gain in subsequent stages. On Days 2 and 3 (reinstatement stage), participants underwent a single 30-sec reinstatement trial on each day, with the reinstatement manipulation varying across groups (see below for details). On Day 4 (test stage), all participants performed the same task as in the training stage again. Performance gain was defined as the subtraction of the baseline performance in the last training trial from performance in the first trial of the test stage.

To dissociate the effects of voluntary movement, somatosensation, and other factors including visual information, we employed three types of manipulation during the reinstatement stage. In an exoskeleton-based reinstatement group (N = 15), a hand-exoskeleton robot (Furuya et al. 2025) reproduced each participant’s learned finger movement recorded during the training stage for 30 sec with the replay of the visual display recorded during the training stage (Fig. 1a; see Methods for details). During this exoskeleton-based reinstatement, participants were instructed to remain relaxed and not move voluntarily. In a voluntary reinstatement group (N = 15), participants were asked to voluntarily execute the same task as in the training stage for one 30-sec trial. In a visual-only reinstatement group (N = 13), only the visual display recorded during the training stage was replayed for 30 sec without any finger movements.

As detailed below, the comparisons of performance gains among the exoskeleton-based, voluntary, and visual-only reinstatement groups would indicate whether the reinstatement of somatosensory information is sufficient to yield performance gain. First, the voluntary reinstatement is expected to result in performance gain according to a prior study (Herszage, Sharon, and Censor 2021). Thus, comparison of performance gains between the exoskeleton-based and voluntary reinstatement groups would show whether performance gain due to the exoskeleton-based reinstatement is comparable to that due to voluntary execution of the trained task. Second, another comparison of performance gains between the exoskeleton-based and visual-only reinstatement groups would reveal whether performance gain shown by the exoskeleton-based reinstatement group occurs due to the reinstatement of somatosensory information. This is because the critical difference between the two groups was the presence of exoskeleton-driven finger movements and their associated somatosensory information. Thus, if the exoskeleton-based reinstatement produces a larger performance gain than the visual-only reinstatement, this demonstrates that somatosensory information leads to performance gain without voluntary execution of the trained task.

Fig. 1c shows performance gains in the exoskeleton-based, voluntary, and visual-only reinstatement groups (see Fig. S1a-c for performance at all stages in each of the three groups). To test if performance gains occur in the three groups, we applied a two-way mixed-model ANOVA on performance with Group (exoskeleton-based, voluntary, vs visual-only reinstatement) as the between-participant factor and Stage (baseline vs test) as the within-participant factor. The results of the ANOVA showed a significant main effect of Stage (F_1,40_ = 43.6044, P < 10^-4^, partial η^2^ = 0.5216, 95% CI of partial η^2^ = [0.3502–0.6235]) and significant interaction between the factors (F_2,40_ = 4.5645, P = 0.0164, partial η^2^ = 0.1858, 95% CI of partial η^2^ = [0.0521–0.3088]), but no significant main effect of Group (F_2,40_ = 0.0773, P = 0.9258, partial η^2^ = 0.0038, 95% CI of partial η^2^ = [0.0000–0.0085]). Subsequently, we found significant simple effects of Stage for the exoskeleton-based (F_1,14_ = 17.5335, P = 0.0009, partial η^2^ = 0.5560, 95% CI of partial η^2^ = [0.2861–0.7342]) and voluntary (F_1,14_ = 25.7381, P = 0.0002, partial η^2^ = 0.6477, 95% CI of partial η^2^ = [0.4608– 0.7601]) reinstatement groups. No significant simple effect of Stage was observed for the visual-only group (F_1,12_ = 4.3333, P = 0.0594, partial η^2^ = 0.2653, 95% CI of partial η^2^ = [0.0017–0.6410]). As the significant interaction between the factors was found, there should be a significant difference in performance gains across the groups. Indeed, performance gains were significantly larger in the exoskeleton-based (two-sample t-test with Shaffer’s modified sequentially rejective Bonferroni procedure; t_40_ = 2.4819, corrected P = 0.0244, Cohen’s d = 0.9784, 95% CI of performance gain difference = [0.6285–4.8382]) and voluntary (t_40_ = 2.7846, corrected P = 0.0244, Cohen’s d = 1.1960, 95% CI of performance gain difference = [1.1310–5.0023]) reinstatement groups, compared to the visual-only reinstatement group. No significant difference in performance gain was observed between the exoskeleton-based and voluntary reinstatement groups (t_40_ = 0.3141, corrected P = 0.7551, Cohen’s d = 0.1015, 95% CI of performance gain difference = [-2.1238–2.7905]). These results suggest that reinstatement of somatosensory information from the learned finger movement is sufficient to produce performance gain comparable to that produced by voluntary execution of the learned movement.

To address alternative possibilities, we conducted a series of control analyses and experiments. First, the baseline performance on the last training trial did not significantly differ across the groups (one-way ANOVA; F_2,40_ = 0.5099, P = 0.6044, partial η^2^ = 0.0249, 95% CI of partial η^2^ = [0.0000–0.1411]; Fig. S1a-c). Thus, group differences in performance gains were not attributable to incidental differences of the baseline performance. Second, the performance gain differences across the groups could not be explained by a speed-accuracy trade-off. A two-way mixed-model ANOVA on error rate (Fig. S2; see Methods for calculation details) revealed a significant main effect of Stage (F_1,40_ = 5.2240, P = 0.0277, partial η^2^ = 0.1155, 95% CI of partial η^2^ = [0.0038–0.2876]), indicating decreased error rates in the test stage. On the other hand, we found no significant main effect of Group (F_2,40_ = 0.2059, P = 0.8148, partial η^2^ = 0.0102, 95% CI of partial η^2^ = [0.0000–0.0360]) or significant interaction between the factors (F_2,40_ = 1.0873, P = 0.3469, partial η^2^ = 0.0516, 95% CI of partial η^2^ = [0.0004–0.1639]). Thus, there were no systematic changes in error rate that could account for the performance gain differences across the groups. Third, for the voluntary reinstatement group no significant performance gain was found in the first day of the reinstatement stage compared to the baseline (Fig. S1b; paired t-test; t_14_ = 0.4847, P = 0.6354, Cohen’s d = 0.1251, 95% CI of performance gain = [-1.1418–1.8084]). This argues against the possibility that performance gains had already emerged before the reinstatement manipulation was applied. The same logic applies to the exoskeleton-based reinstatement group since this group followed the same procedure until the beginning of the first day of the reinstatement stage. Fourth, we conducted Control Experiment 1 (N = 12) which omitted the reinstatement stage (see Methods for details). Previous research on visual skill learning showed that performance can continue to improve for several days after initial training without additional training (Stickgold, James, and Hobson 2000). This raises a possibility that performance gain might have accumulated over several days after the training stage, irrespective of the reinstatement manipulations. However, this control experiment revealed no significant performance gain at the test stage, compared to the baseline (Fig. S3; t_11_ = 0.5195, P = 0.6137, Cohen’s d = 0.1500, 95% CI of performance gain = [-2.6973–4.3640]), arguing against the possibility. Finally, we conducted Control Experiment 2 (N = 14) in which participants were exposed to exoskeleton-driven sequential finger movements unrelated to the trained sequence during the reinstatement stage. This experiment tested the possibility that exposure to exoskeleton-driven finger movement led to general factors such as enhanced arousal and warm-up effects, yielding the performance gain for the exoskeleton-based reinstatement group. However, we found no significant performance gain (Fig. S4; paired t-test; t_13_ = 0.4288, P = 0.6751, Cohen’s d = 0.1146, 95% CI of performance gain = [-2.0193–3.0193]). This is inconsistent with the general facilitation account and instead suggests that a certain degree of sequence compatibility with the trained sequence is required for reinstatement-driven performance gain.

Together, the results of Experiment 1 suggested that brief exoskeleton-based reinstatement of training-related somatosensory information is sufficient to strengthen a motor memory of the learned sequential skill without voluntary movement.

### Exoskeleton-based reinstatement strengthens the skill even under high cognitive load

Was the performance gain observed in Experiment 1 driven by afferent somatosensory information itself, or was it produced by cognitive processes such as mental rehearsal? One may argue that the exoskeleton-driven reproduction of the trained finger movement could have served as cues for covert mental rehearsal of the trained movement, which might have resulted in performance gain. If this possibility is true, performance gain should not be observed when participants perform a demanding task that hinders access to the exoskeleton-driven finger movement and therefore minimizes the opportunity for mental rehearsal during the reinstatement stage. By contrast, if performance gain was driven by afferent somatosensory information itself, the gain should still be observed after the reinstatement stage with the demanding task.

Experiment 2 with a new group of participants (N = 13) was designed to dissociate the contribution of afferent somatosensory information from that of mental rehearsal (see Methods for details). Participants were trained and tested in the same way as the exoskeleton-based reinstatement group in Experiment 1, with one exception. During the reinstatement stage, while participants were exposed to the exoskeleton-driven finger movements, they simultaneously performed a demanding two-back working memory (WM) task on a sequence of object images presented on the display (Fig. 2a).

**Fig. 2.**
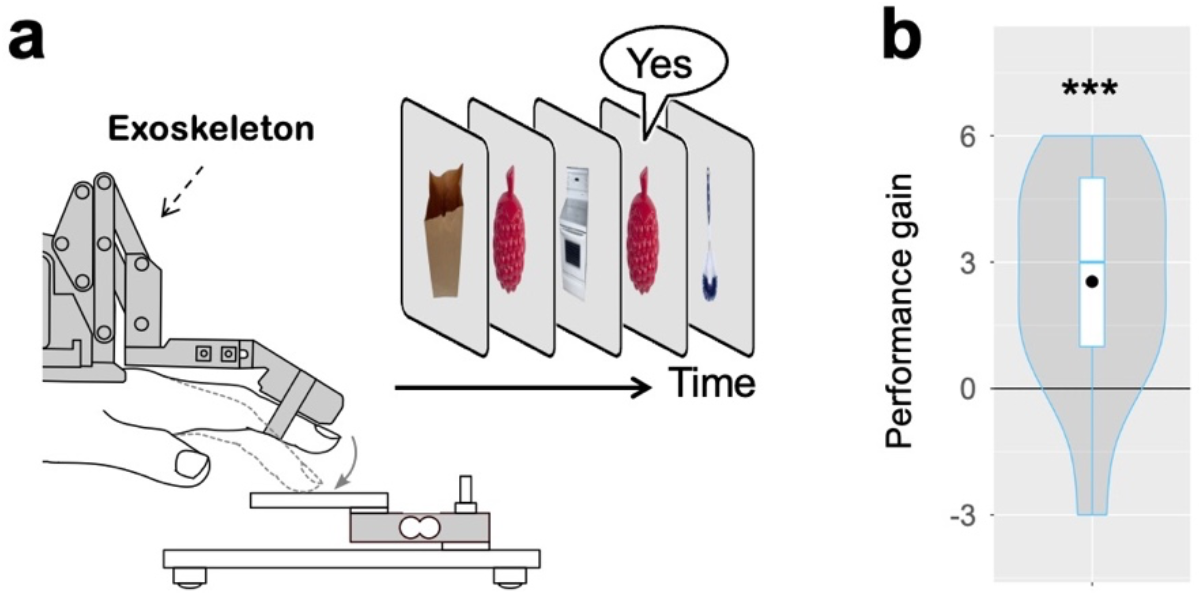
Procedures and results of Experiment 2 **a**, Two-back working memory (WM) task. In addition to the exposure to the exoskeleton-driven finger movements, images of objects were sequentially presented on a display. Participants were asked to respond “yes” verbally when the current image matched the one presented two images earlier. **b**, Performance gain in the test stage, relative to the baseline (N = 13). Box plots are overlaid on violin plots. Black dots indicate mean values across participants. Violin plots show kernel probability densities of individual values. ***P < 0.005.

Despite the concurrent WM task during the reinstatement stage, participants showed a significant performance gain from the baseline to the test stage (Fig. 2b; paired t-test; t_12_ = 3.6078, P = 0.0036, Cohen’s d = 1.0006, 95% CI of performance gain = [1.0054– 4.0715]; see Fig. S1d for performance at all stages). This result further supports the view that afferent somatosensory information is sufficient to drive later performance gain.

However, this interpretation assumes that the WM task effectively limited access to sequential information driven by the exoskeleton. To confirm this assumption, we conducted Control Experiment 3 (N = 13) based on a dual-task paradigm (Fig. S5a; see Methods for details). In single-task conditions, they were asked to perform either a finger discrimination task or the same visual two-back WM task as in Experiment 2. In the finger discrimination task, exoskeleton-driven sequential finger movement was followed by a two-alternative forced-choice (2AFC) test, in which the presented sequential movement and a lure sequential movement were presented in succession. Participants were asked to indicate which sequential movement matched the one presented earlier. In the dual-task condition, they were asked to concurrently perform the two tasks while being instructed to prioritize the WM task. Fig S5b shows percent reduction of performance for each task in the dual-task condition compared to the single-task condition. Performance reduction in the finger discrimination task was significantly greater than that in the WM task (paired t-test; t_12_ = 6.2560, P < 10^-4^, Cohen’s d = 1.7351, 95% CI of performance reduction = [-41.8419 – -20.2253]%), indicating that finger-discrimination performance declined close to the chance level in the dual task condition (Fig. S5c; see Fig. S5d for WM performance). Thus, participants could not reliably access the trained sequence while concurrently performing the demanding WM task. These results suggest that participants were unable to perform covert mental rehearsal of the trained finger movements during the reinstatement stage of Experiment 2.

### Passive exposure to a partially matched sequence strengthens the skill

In Experiments 1 and 2, exoskeleton-based reinstatement reproduced somatosensory information derived from the trained sequence. These results raise a mechanistic question about the format in which an acquired motor memory can be accessed by afferent somatosensory information. One possibility is that the reinstatement of somatosensory information modifies motor memory in a strictly content-specific manner. Under this account, the reinstatement of somatosensory information would have to specify the precise motor memory to be strengthened, and therefore the identical trained sequence should be required. An alternative possibility is that the reinstatement of somatosensory information serves as an access cue that makes a related motor memory eligible for strengthening. Under this account, the reinstated sequence need not match the entire trained sequence. Instead, it may be sufficient for the sequence to share part of its transition structure with the trained sequence, because such partial overlap could provide sufficient access to the relevant motor memory.

Experiment 3 with a new group of participants (N = 14) was designed to distinguish between these possibilities by testing whether passive exoskeleton-based exposure to a partially matched sequence during the reinstatement stage can still produce a performance gain. The total variation distance between the 2-gram frequency distributions of the partially matched and trained sequences was 0.64 (see Methods for details). All other procedures, including the concurrent WM task, were identical to those in Experiment 2. We found a significant performance gain from the baseline to the test stage (Fig. 3; paired t-test; t_13_ = 3.5143, P = 0.0038, Cohen’s d = 0.9392, 95% CI of performance gain = [1.0732–4.4982]; see Fig. S1e for performance at all stages). The results of Experiment 3 suggest that partially matched somatosensory information is sufficient to drive the process that strengthens motor memory.

**Fig. 3.**
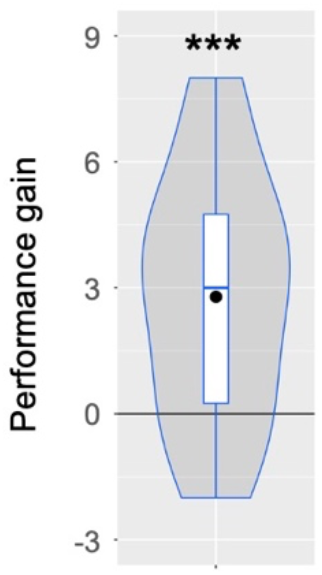
Results of Experiment 3 Performance gain in the test stage, relative to the baseline (N = 14). Box plots are overlaid on violin plots. Black dots indicate mean values across participants. Violin plots show kernel probability densities of individual values. ***P < 0.005.

### Voluntary reinstatement requires stricter sequence specificity than exoskeleton-based reinstatement

In Experiment 1, exoskeleton-based reinstatement and voluntary reinstatement produced comparable performance gains when the reinstated sequence was identical to the trained sequence (Fig. 1c). However, this behavioral equivalence does not establish that afferent somatosensory information and voluntary motor execution access motor memory under the same constraints. Experiment 3 showed that passive exoskeleton-based exposure to a partially matched sequence was sufficient to yield performance gain, suggesting that the reinstatement of somatosensory information can access the trained memory through partial transition structure rather than requiring the complete trained sequence. We therefore asked whether voluntary reinstatement has the same tolerance for partial sequence overlap. If voluntary reinstatement relies on the same mechanism, voluntary execution of the same partially matched sequence should also induce performance gain. If, instead, voluntary execution imposes a stricter specificity constraint on the memory to be accessed, then executing the partially matched sequence should not yield performance gain.

Experiment 4 (N = 17) tested this prediction by having a new group of participants voluntarily execute the same partially matched sequence used in Experiment 3 during the reinstatement stage (Fig. 4a; see Methods for details). Unlike the passive exposure in Experiment 3, the voluntary execution of the partially-matched sequence did not yield a significant performance gain at the test stage (Fig. 4b; paired t-test; t_16_ = 0.9330, P = 0.3647, Cohen’s d = 0.2263, 95% CI of performance gain = [-0.7483–1.9248]; see Fig. S1f for performance at all stages). Moreover, the performance gain in Experiment 3 was significantly higher than that in this Experiment 4 (two-sample t-test; t_29_ = 2.1696, P = 0.0393, Cohen’s d = 0.7934, 95% CI of performance gain difference = [0.1161–4.2788]). These results suggest that the performance gains produced by exoskeleton-based reinstatement and voluntary reinstatement are based on different mechanisms. Specifically, somatosensory information may act as an access cue and broadly drive the modification of related motor memories. On the other hand, voluntary motor execution appears to impose a specificity such that only a motor skill memory related to the executed sequence is eligible for strengthening.

**Fig. 4.**
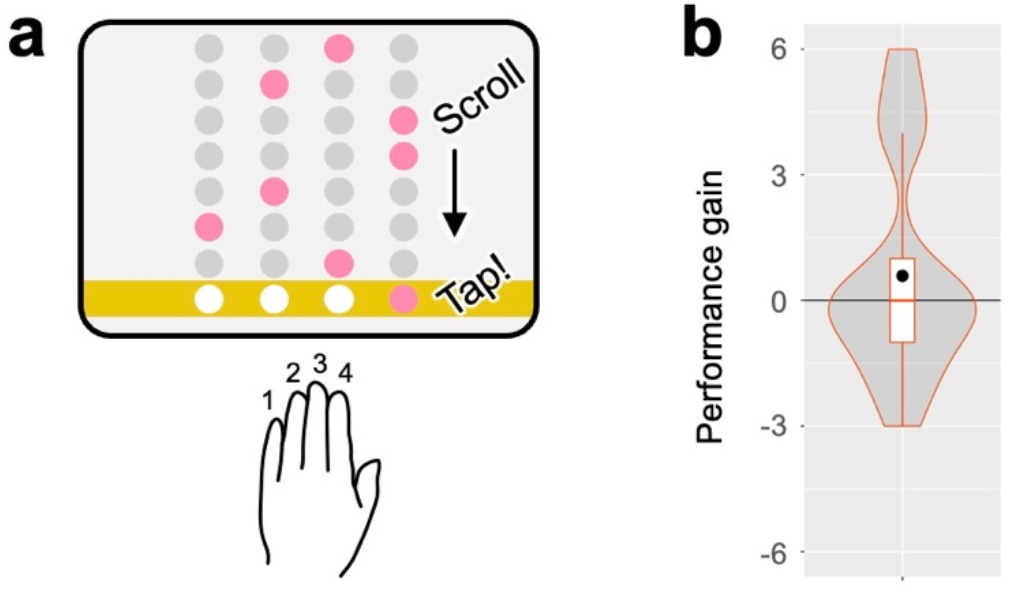
Procedures and results of Experiment 4 **a**, Voluntary execution of the partially-matched sequence (see Methods for details). Four circles on the yellow line indicated the current target finger, and circles above the yellow line indicated the upcoming target fingers. A metronome sound was presented, with the interval adjusted to match the tapping speed during the training stage for each participant. Participants were asked to tap in synchrony with the metronome sound. **b**, Performance gain in the test stage, relative to the baseline (N = 17). Box plots are overlaid on violin plots. Black dots indicate mean values across participants. Violin plots show kernel probability densities of individual values.

## Discussion

The present study shows that afferent somatosensory information can drive the modification of an already acquired motor skill memory. Motor memory modification has typically been studied in the context of voluntary movement, where motor commands, sensory predictions, performance feedback, and somatosensation are tightly coupled (Wolpert, Ghahramani, and Jordan 1995; Todorov and Jordan 2002; Shadmehr, Smith, and Krakauer 2010). Within this framework, somatosensory information has therefore often been assigned a supportive role such as feedback for error correction or calibration of sensorimotor mappings. By isolating afferent somatosensory information from voluntary motor execution, the present study provides evidence against this restricted view. Brief, 30-sec exoskeleton-based reinstatement of the learned finger movements produced subsequent performance gain comparable to that induced by voluntary execution of the trained sequence (Fig. 1c), whereas no significant gain was detected after visual-only reinstatement. Thus, somatosensory information is not merely a consequence of movement that supports learning; it can itself serve as an effective input for modifying an existing motor skill memory.

Several alternative explanations are unlikely to account for this effect. First, the performance gain induced by exoskeleton-based reinstatement was preserved even when participants performed a demanding working memory task during reinstatement (Fig. 2b). This finding argues against the possibility that the gain was primarily driven by covert rehearsal or mental simulation of the trained sequence. Second, visual-only reinstatement did not induce performance gain, indicating that the effect cannot be attributed to generic exposure to task-related visual cues or environments (Fig. 1c). Third, the control experiments made it unlikely that the performance gain reflected nonspecific factors such as enhanced arousal, warm-up, tactile stimulation, or repeated mounting of the exoskeleton (Fig. S4a). Together, these findings support the conclusion that task-relevant afferent somatosensory information was sufficient to trigger later improvement in the trained motor skill.

Our findings clarify what somatosensory information can do in motor memory modification. Prior studies have shown that somatosensory representations change with motor learning and are associated with motor memory retention (Ostry et al. 2010; Hirano, Sakurada, and Furuya 2020), and that disrupting somatosensory cortical processing can impair the retention or consolidation of newly learned motor memory (Wali 2020; Ebrahimi, van der Voort, and Ostry 2024). Passive movement studies further suggest that somatosensory information can contribute to the recognition of previously learned movements (Kumar, van Vugt, and Ostry 2021). However, these studies did not determine whether afferent somatosensory information, isolated from voluntary motor execution, can itself modify an already acquired motor skill memory. The present results fill this gap by suggesting that somatosensory information can act as an access signal for an existing motor memory. That is, reproducing the somatosensation of a learned movement may reinstate a sensorimotor state associated with the original training episode, thereby triggering a process that further strengthens the corresponding motor memory. In this sense, somatosensory reinstatement does not simply provide online correction signals for current performance; rather, it can initiate offline modification of a previously acquired skill memory even in the absence of overt motor performance.

The present results also suggest that somatosensory-based and voluntary movement-based routes to motor memory modification have different functional constraints. This idea is consistent with the broader view that a motor memory is not a unitary trace, but consists of partially dissociable components that can undergo distinct forms of offline processing (Cohen et al. 2005; Robertson 2009). When the reinstated sequence exactly matched the trained sequence, exoskeleton-based reinstatement and voluntary reinstatement produced comparable performance gains (Fig. 1c). However, this equivalence did not generalize to the partially matched sequence. Passive exoskeleton-based exposure to the partially matched sequence still improved later performance of the trained sequence (Fig. 3), whereas voluntary execution of the same partially matched sequence did not (Fig. 4b). This dissociation suggests that afferent somatosensory information can engage motor memories through partially overlapping sensorimotor structure, whereas voluntary execution appears to impose a stricter selection constraint on which memory is strengthened. Put differently, somatosensory reinstatement may provide relatively broad access to related motor memories, whereas voluntary execution may preferentially modify the memory corresponding to the executed action.

The present study builds on the motor memory reactivation and consolidation framework established in prior behavioral studies. A central idea in this framework is that motor memories continue to be processed offline after training, and that reactivation can render an existing memory susceptible to subsequent modification (Robertson 2009, 2012). Consistent with this view, brief reactivation of an existing motor memory can make that memory modifiable (Censor, Dimyan, and Cohen 2010), prevent interference (Herszage and Censor 2017), or induce subsequent performance gains (Herszage, Sharon, and Censor 2021). Here, we use the term reinstatement in this behavioral sense, referring to an experimentally controlled procedure that reinstates information associated with the originally learned movement. The key extension is that reinstatement was achieved not by voluntary execution, but by externally reproducing the somatosensation of the learned movement using a hand-exoskeleton robot. Our findings extend the motor memory reactivation framework by identifying afferent somatosensory information as a route through which an existing motor skill memory can be reinstated and further strengthened.

This behavioral notion of reinstatement should be distinguished from endogenous neural replay during offline periods. Previous studies have linked micro offline skill consolidation with replay of neural activity representing a trained motor sequence during waking rest (Bönstrup et al. 2019; Buch et al. 2021; Bönstrup et al. 2020). The present study did not measure neural replay. An important future question is how behaviorally defined reinstatement relates to subsequent neural replay and offline performance gain.

More broadly, the present findings support a view in which bodily input plays a more active role across multiple stages of motor learning than previously thought. Previous studies have shown that somatosensory information can facilitate initial motor skill acquisition, support recognition of previously experienced movements, and enhance motor performance under specific conditions (Bernardi, Darainy, and Ostry 2015; Kumar, van Vugt, and Ostry 2021; Furuya et al. 2025). Although in a different sensory modality and learning domain, recent work in visual perceptual learning has similarly shown that post-training sensory stimulation can modulate wakeful consolidation and alter subsequent performance (Marzoll et al. 2022; Yang et al. 2025). The present study extends this broader principle to motor memory by showing that brief reinstatement of training-related afferent input can further strengthen an already acquired sequential motor memory without voluntary execution. Together, these findings suggest that somatosensation is not merely a downstream consequence of motor commands, but can serve as an active input that shapes motor learning from initial acquisition to post-acquisition memory strengthening.

In summary, the present study resolves a key controversy in the role of somatosensory information in motor memory modification. Somatosensory information is not limited to a supportive role in feedback or calibration. When isolated from voluntary motor execution, brief reinstatement of training-related afferent somatosensory information can drive subsequent improvement in a learned motor skill. The results further suggest that this somatosensory route differs from voluntary movement-based modification in its specificity constraints, providing a route by which relevant motor memories can be accessed and strengthened without overt action.

## Methods

### Participants

A total of 176 naïve participants (right-handed, no history of neurological disorders) were enrolled in this study. The experimental protocols were approved by the Institutional Review Board. Prior to participation, all participants provided written informed consent. Data from 50 participants were subsequently excluded from the analyses (see Exclusion of participants for details). Consequently, the final analyses included data from 126 participants (18 to 32 years old; 100 males and 26 females).

We recruited participants so that the sample size would fall approximately within the same range (N = 10–15) per group as in prior studies using the same sequential finger movement task with analogous designs (Censor, Dimyan, and Cohen 2010; Herszage and Censor 2017; Herszage, Sharon, and Censor 2021).

### Sequential finger movement task

Participants performed a sequential finger movement task (Fig. 1a) in which they repeatedly tapped a 5-item sequence (4–1–3–2–4) shown on a display as quickly and accurately as possible (Bönstrup et al. 2019, 2020; Buch et al. 2021; Censor, Dimyan, and Cohen 2010; Herszage and Censor 2017; Herszage, Sharon, and Censor 2021). Participants placed their left-hand fingers on 5-kg load cells that detected finger taps. Taps 1–4 corresponded to the little, ring, middle, and index fingers, respectively. Each tap produced an asterisk on the display, which accumulated from left to right as the trial progressed. Each trial lasted 30 sec and was followed by a 30-sec rest period.

Performance and its gain were defined as follows. Each correct completion of the full sequence was counted as one successful sequence. Performance was defined as the number of correctly completed sequences in a 30-sec trial, following a prior study (Herszage, Sharon, and Censor 2021). Performance gain was calculated as performance in the first test trial minus performance in the last training trial. An error rate for each trial was calculated as (the total number of taps – 5 × performance) / the total number of taps.

### Experiment 1

The objective of Experiment 1 was to test whether the reinstatement of training-related somatosensory information is sufficient to improve performance in the sequential finger movement task without voluntary movement. Experiment 1 consisted of a 1-day training stage, a 2-day reinstatement stage, and a 1-day test stage in this order (Fig. 1b), following a prior study (Herszage, Sharon, and Censor 2021). The experiment was conducted every other day. During the training stage, all participants performed 12 trials of the sequential finger movement task. During the reinstatement stage, participants underwent a single 30-sec reinstatement trial on each day. The manipulations varied across groups (see below for details). During the test stage, all participants performed the same sequential finger movement task for five trials. To avoid additional training effect, only the first trial was used to calculate performance gains. Note that all participants, regardless of group, wore the data glove on their left hand during the training and test stages. Similarly, the hand-exoskeleton robot was mounted on their left hand during the reinstatement stage. It is thus unlikely that the differences in performance gain across the groups were due to differences in the equipment (i.e., the data glove and hand-exoskeleton robot).

Participants were randomly assigned to three groups: exoskeleton-based (N = 15), voluntary (N = 15), and visual-only (N = 13) reinstatement groups. In the exoskeleton-based reinstatement group, the hand-exoskeleton robot (Furuya et al. 2025) was used to reproduce the finger movements performed by each participant during the training stage (Fig. 1a). For each participant, the trial with the best performance during the training stage was selected, and the voluntary finger movements in the selected trial were used to generate exoskeleton-driven finger movements for the reinstatement stage (see Generating exoskeleton-driven finger movement for details). The 5-item sequence (4–1–3–2–4) was shown on the display, and asterisks accumulated in synchrony with the exoskeleton-driven finger movements. Participants in this group were asked to stay relaxed and not to move voluntarily during the reinstatement stage. In the voluntary reinstatement group, participants performed the trained finger movement task. In the visual-only reinstatement group, only the visual stimuli recorded during the training stage (5-item sequence and accumulating asterisks) were replayed while the exoskeleton robot locked the finger joints.

### Experiment 2

The objective of Experiment 2 with a new group of participants (N = 13) was to test whether performance gain due to the exoskeleton-based reinstatement occurs when participants were engaged in a demanding task. The procedures were identical to those for the exoskeleton-based reinstatement group of Experiment 1, except that participants performed a two-back working memory (WM) task (Fig. 2a), in addition to the exposure to exoskeleton-driven finger movements. In this task, images of everyday objects (e.g., a refrigerator, a doll) were sequentially presented on a display, each for 0.2 sec followed by a 0.7 sec inter-stimulus interval. Participants were instructed to respond “yes” verbally when the current image matched the one presented two images earlier. A total of 45 images were presented, of which 25% were target (“yes”) trials. During the task, the exoskeleton robot reproduced the finger movements that participants had performed during the training stage, starting after a variable delay from trial onset.

### Experiment 3

The objective of Experiment 3 with a new group of participants (N = 14) was to test whether performance gain occurs as a result of exposure to a sequence that partially matches the trained sequence during the reinstatement stage. The procedures were identical to those in Experiment 2, except that a partially matched sequence was used instead of the trained sequence during the reinstatement stage. This sequence was generated by shuffling a concatenated repetition of the trained sequence (4–1–3–2–4), with the constraint that no three consecutive 4s occurred, thereby preserving the marginal frequency of finger movements and allowing some trained transitions to occur while disrupting the full trained transition structure. The total variation distance between the 2-gram frequency distributions of the partially matched and trained sequences was 0.64. The timing of the exoskeleton-driven movements was adjusted for each participant so that the mean inter-movement interval matched the mean inter-tap interval during the training stage.

### Experiment 4

The objective of Experiment 4 with a new group of participants (N = 17) was to test whether performance gain occurs as a result of voluntary execution of sequential movements that partially match the trained sequence during the reinstatement stage. The procedures were identical to those used for the voluntary reinstatement group in Experiment 1, except that participants tapped the same partially matched sequence used in Experiment 3. As shown in Fig. 4a, circles and a yellow line were displayed. Four circles on the yellow line indicated the current target finger, and circles above the yellow line indicated the upcoming target fingers. A metronome sound was presented, with the interval adjusted to match the tapping speed during the training stage for each participant. Participants were instructed to tap in synchrony with the metronome sound. Once the correct tap was made, the sequence scrolled downward, bringing the next target onto the yellow line.

### Control Experiment 1

The objective of Control Experiment 1 with a new group of participants (N = 12) was to test whether performance gain occurs over several days after the training stage. The procedures were identical to those in Experiment 1, except that the reinstatement stage was omitted. As in Experiment 1, there was a 5-day interval between the training and test stages.

### Control Experiment 2

The objective of Control Experiment 2 with a new group of participants (N = 14) was to test whether performance gain occurs as a result of exposure to sequential finger movement unrelated to the trained sequence during the reinstatement stage. The procedures were identical to those for the passive reinstatement group in Experiment 1. The only exception was that a new sequence was used during the reinstatement stage. The new sequence (4– 2–1–4–3) matched the trained sequence (4–1–3–2–4) in the numbers of movements for each finger. However, unlike the partially matched sequence used in Experiments 3 and 4, this new sequence contained one of the trained first-order transitions (4–1, 1–3, 3–2, 2–4, 4–4).

### Control Experiment 3

The objective of Control Experiment 3 with a new group of participants (N = 13) was to test whether the WM task used in Experiment 2 effectively limits access to sequential information in the exoskeleton-driven finger movements (Fig. S5a). A finger discrimination task was combined with the WM task used in Experiment 2. In the finger discrimination task, one sequence was selected from a set consisting of the veridical sequence (4-1-3-2-4) and all its shuffled variants and was repeatedly presented as the target by the exoskeleton robot for 30 sec. This was followed by a two-alternative forced-choice (2AFC) test, in which the target and a lure sequence (i.e., a non-target sequence from the same set) were presented once in succession. Participants were asked to indicate which sequence matched the target. The experiment consisted of three conditions: finger discrimination only, WM only, and dual-task conditions. In the dual-task condition, participants performed both tasks simultaneously while prioritizing performance on the WM task. Eleven trials were conducted for each condition in a randomized order.

For the finger discrimination task, performance was quantified as the proportion of correct responses in the 2AFC test (Fig. S5c). The performance of the WM task (d’) was assessed with a standard procedure based on hits, misses, false alarms, and correct rejections (Fig. S5d).

### Generating exoskeleton-driven finger movements

Finger angle time series recorded by the data glove during the training stage were low-pass filtered at 10 Hz and segmented into individual cycles of the task sequence (4–1–3–2–4), with movement onset defined by velocity peaks. Each segment was temporally normalized to a common scale (0–100%) via interpolation. Temporal profiles were aligned across cycles using characteristic landmarks (e.g., onset, intermediate points, and offset), together with piecewise temporal warping to match their mean timing across cycles. The mean trajectory was computed across cycles for each finger and then temporally arranged based on these landmark points so that finger movements followed a given sequence order, including sequences not directly observed.

The resulting cycles of the sequence were combined to generate a continuous 30-sec movement sequence, with the number of cycles determining the movement speed. This sequence was reproduced by the exoskeleton robot to deliver passive finger movements.

### Exclusion of participants

We excluded 16 participants who did not show stable skill acquisition during the training stage as the aim of this study was to test differential effects of experimental manipulations in the reinstatement stage on performance gain after the initial acquisition. Specifically, we fit an exponential function to performance time-course across the 12 trials in the training stage and excluded participants whose coefficient of determination for the fit was below 0.5. It has been reported that individual learning curves are typically well captured by an exponential function (Heathcote, Brown, and Mewhort 2000). Consistent with this, exponential fitting is the standard approach to evaluate learning in the sequential finger movement task used in this study (Bönstrup et al. 2019, 2020; Buch et al. 2021).

Additionally, 34 participants were excluded from the analyses for the following reasons: 15 due to equipment malfunction, 6 due to incompatibility with the experimental apparatus (e.g., mismatched joint angles between the hand and the data glove), and 13 due to failure to complete the experiment (e.g., illness or withdrawal).

### Statistics

All the statistical tests performed in this study were two-tailed. The significance threshold was set to an alpha level of 0.05. *t*-tests and ANOVA were performed utilizing the function anovakun in the R statistical software package. Effect size for *t*-tests was quantified using Cohen’s d, while partial η^2^ served as the measure of effect size for ANOVAs. The computation of the 95% confidence intervals was based on a bias-corrected and accelerated (BCa) method over 2,000 bootstrap repetitions. For ANOVAs, we applied Mendoza’s multi-sample sphericity test to evaluate the sphericity assumption. When necessary, we adjusted the degrees of freedom using Greenhouse-Geisser’s ε to correct any violations.

### Apparatus

Experiments were controlled using PsychoPy on Windows OS. Visual stimuli were presented on an LED display operated at a refresh rate of 60 Hz.

To record and reproduce participants’ finger movement in Experiments 1–4, we used a custom-made data glove and hand-exoskeleton robot developed in a prior study (Furuya et al. 2025). In the training stage, we recorded finger angles using the data glove at a temporal resolution of 1000 Hz. To generate exoskeleton-driven finger movements, the exoskeleton robot applied force to the proximal phalanx of each of the four fingers and to flex and extend the MP joint of each finger with an angular resolution of less than 1.0 deg. The exoskeleton robot moved according to a predefined sequence of joint angles of the fingers over time. The detailed control system of the exoskeleton robot and its specifications are shown in the prior study (Furuya et al. 2025).

## Acknowledgements

We express our deepest gratitude to S. Furuya for developing and providing the exoskeleton device and his invaluable support and advice on experimental designs, analyses, and the manuscript. This work was supported by JSPS Kakenhi Grants 24K02848 (to H.O.), 20H05715 (to K.S.), and JST Moonshot R&D Grant JPMJMS2013 (to K.S.).

