## Supplemental Data 1 for "Somatosensory information drives modification of a motor memory"

### Supplementary figures

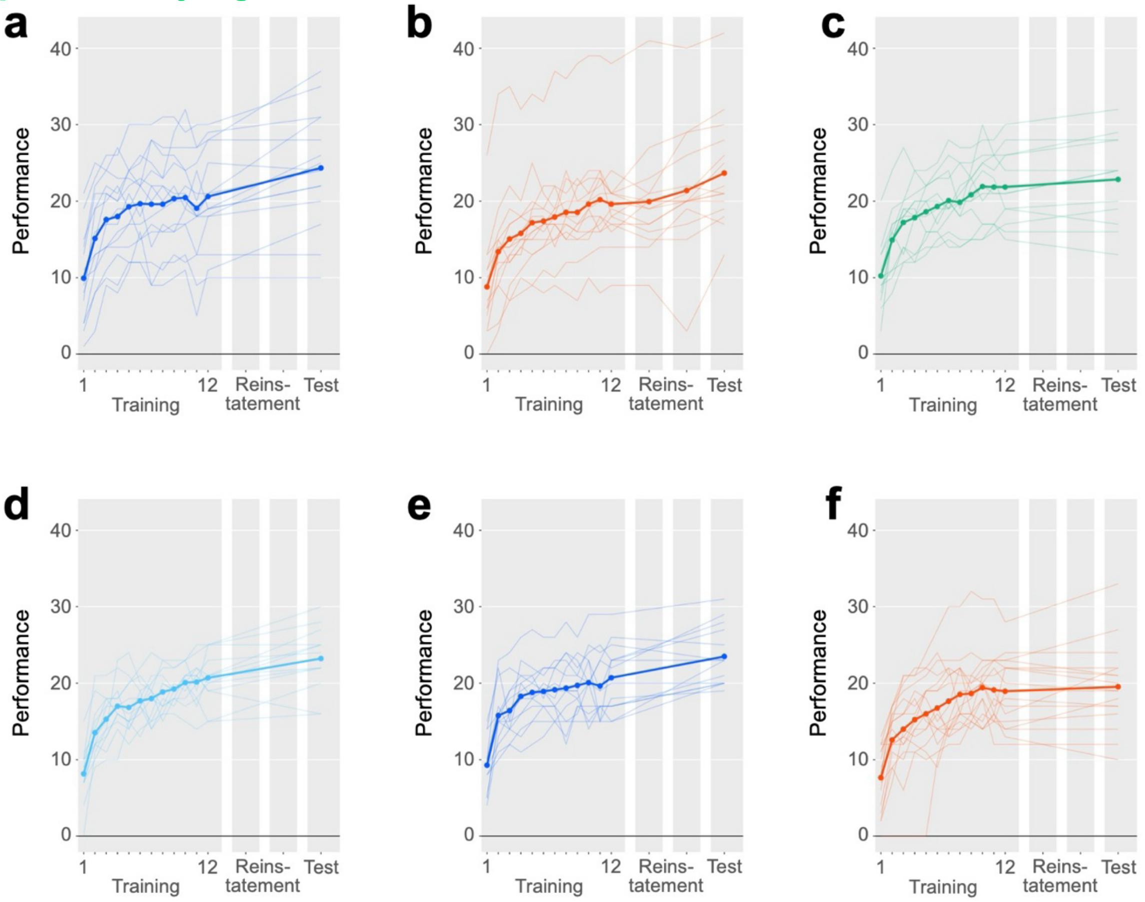


Fig. S1: Performance in the sequential movement task during the training, reinstatement, and test stages of Experiments 1–4

**a**, Performance in the exoskeleton-based reinstatement group of Experiment 1 (N = 15). Thick lines represent mean values across participants, and thin lines indicate individual values. **b**, Performance in the voluntary reinstatement group of Experiment 1 (N = 15). **c**, Performance in the visual-only reinstatement group of Experiment 1 (N = 13). **d**, Performance in Experiment 2 (N = 13). **e**, Performance in Experiment 3 (N = 14). **f**, Performance in Experiment 4 (N = 17).

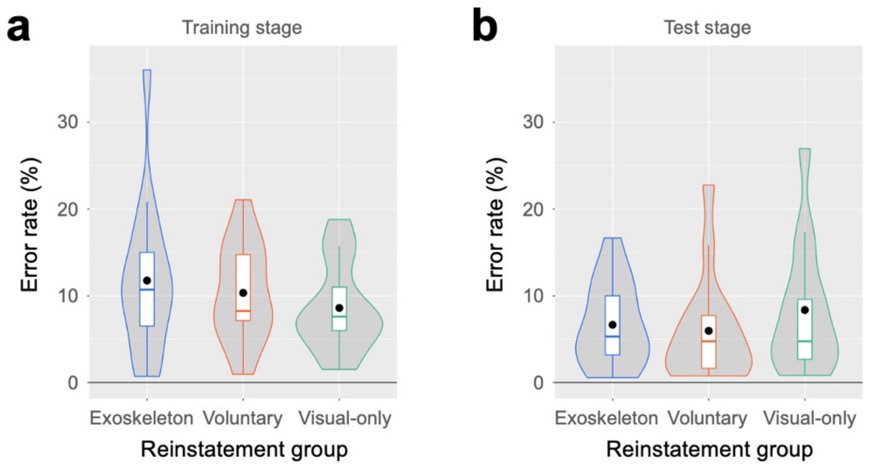


Fig. S2: Error rates in Experiment 1
**a**, Error rates in the training stage for the exoskeleton-based (N = 15), voluntary (N = 15), and visual-only (N = 13) reinstatement groups. Box plots are overlaid on violin plots. Black dots indicate mean values across participants. Violin plots show kernel probability densities of individual values. **b**, Error rates in the test stage.

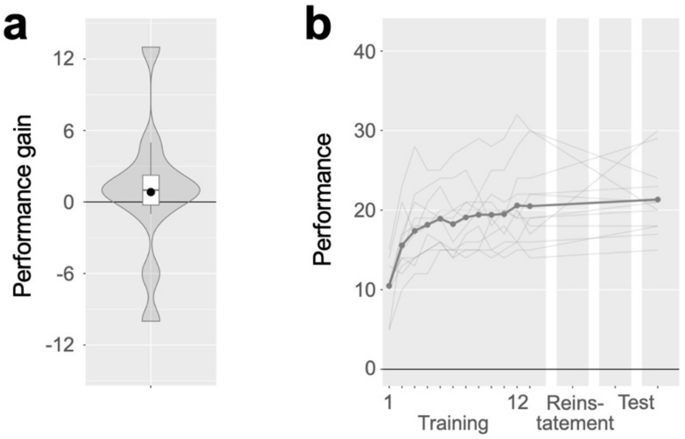


Fig. S3: Results of Control Experiment 1

**a**, Performance gain in the test stage, relative to the baseline (N = 12). Box plots are overlaid on violin plots. Black dots indicate mean values across participants. Violin plots show kernel probability densities of individual values. **b**, Performance in the training and test stages. Thick lines represent mean values across participants, and thin lines indicate individual values.

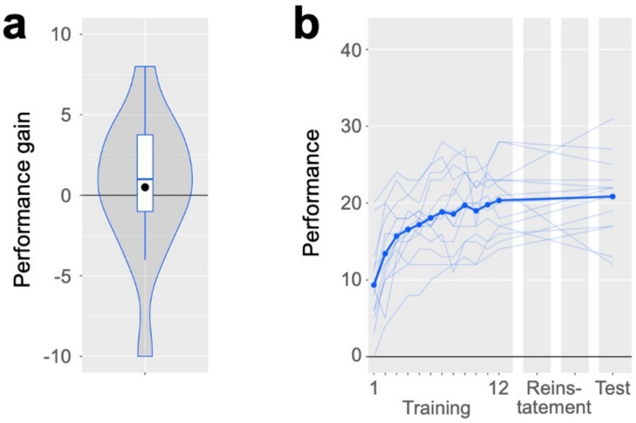


Fig. S4: Results of Control Experiment 2

**a**, Performance gain in the test stage, relative to the baseline (N = 14). Box plots are overlaid on violin plots. Black dots indicate mean values across participants. Violin plots show kernel probability densities of individual values. **b**, Performance in the training and test stages. Thick lines represent mean values across participants, and thin lines indicate individual values.

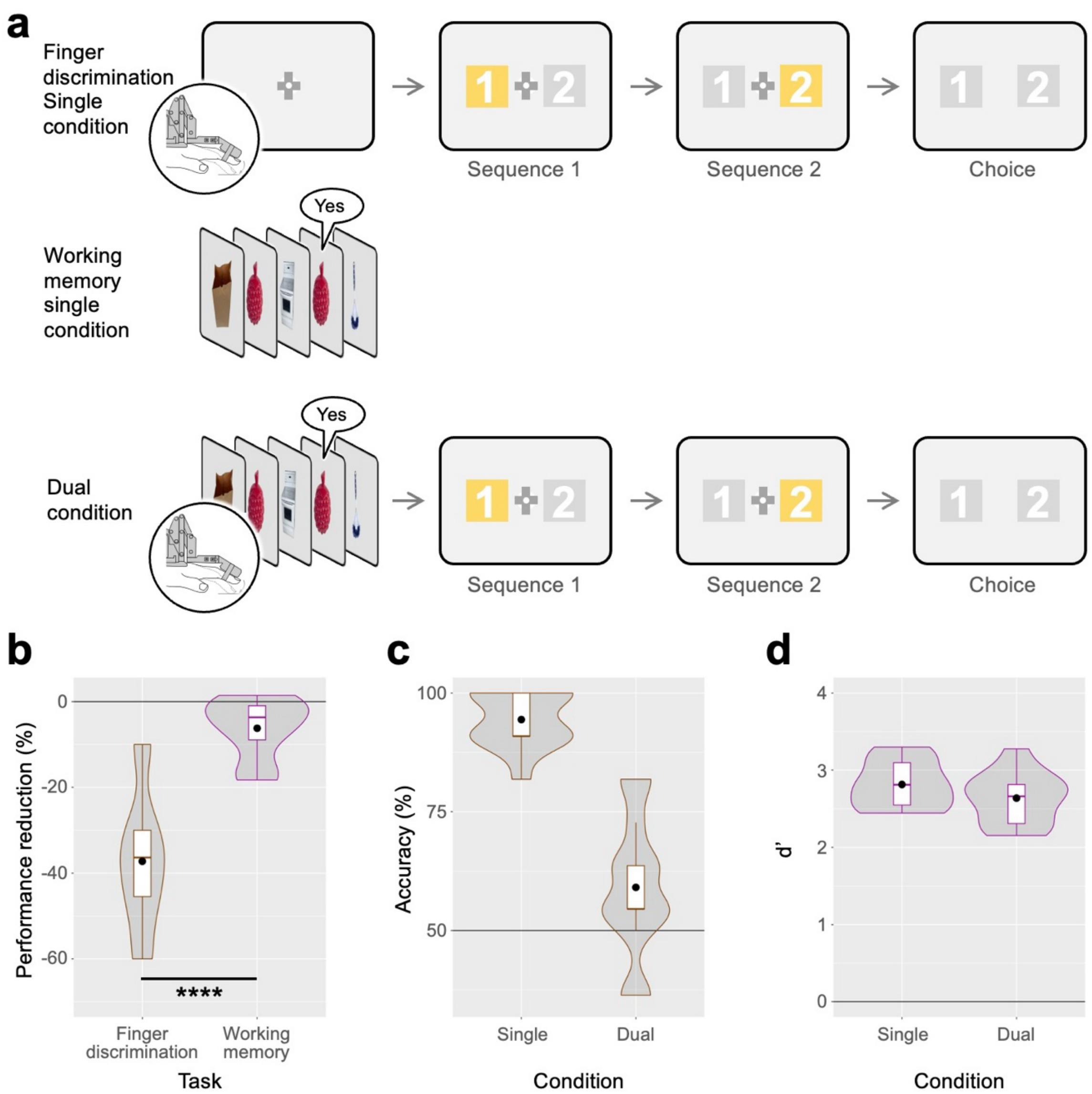


Fig. S5: Procedures and results of Control Experiment 3

**a**, Top: single-task condition with a finger discrimination task. In this task, exoskeleton-driven sequential finger movement was followed by a two-alternative forced-choice (2AFC) test, in which the presented sequential movement and a lure sequential movement were presented in succession. Participants were asked to indicate which sequence matched the one presented earlier. Middle: single-task condition with the same working-memory (WM) task in Experiment 2. Bottom: dual-task condition. In this condition, participants were asked to concurrently perform the two tasks while being instructed to prioritize the WM task. **b**, Percent reduction of performance in the dual-task condition relative to the single-task condition for each task (N = 13). Box plots are overlaid on violin plots. Black dots indicate mean values across participants. Violin plots show kernel probability densities of individual values. ****P < 0.001. **c**, Performance in the finger discrimination task in the single- and dual-task conditions. **d**, Performance in the WM task in the single- and dual-task conditions.
